# White spot syndrome virus endogenous viral elements (EVE) revealed by circular viral copy DNA (cvcDNA) in shrimp

**DOI:** 10.1101/2023.08.30.554989

**Authors:** Suparat Taengchaiyaphum, Jiraporn Srisala, Prapatsorn Wongkhaluang, Suchitraporn Sukthaworn, Joaquin Macias, Iman Ihsan Udin, Madhu Babu Chokkara, Mohammed Musthafa Athikkavil, Victoria Alday-Sanz, Timothy W. Flegel, Kallaya Sritunyalucksana

## Abstract

Circular, viral copy DNA (cvcDNA) can reveal the existence of endogenous viral elements (EVE) in shrimp genomic DNA. Here we describe isolation and sequencing of cvcDNA from a breeding stock of the whiteleg shrimp *Penaeus vannamei*. The stock was developed by onward breeding and selection for white spot syndrome virus (WSSV)-free individual survivors of white spot disease outbreaks. The stock exhibits high tolerance to WSSV. A pool of DNA extracted from 10 shrimp from this stock was subjected to cvcDNA isolation and amplification followed by high throughput sequencing. This revealed DNA fragments corresponding to locations covering much of the 300,000 bp WSSV genome. However, high frequency-read-fragments (HFRF) mapped to a surprisingly small region of approximately 1,400 bp. We hypothesized that the HFRF reflected their selection due to provision of tolerance to WSSV. Four PCR primer sets were designed to cover the 1,365 bp mapped region. One pair (Set 1) targeted the whole 1,365 bp mapped region, while another 3 primer sets (Set 2-4), targeted regions within the 1,365 bp target. All 4 primer sets were used with DNA samples from each of 36 shrimp from the same breeding stock (including the 10 used for cvcDNA preparation). Individual positive PCR results varied widely from shrimp to shrimp, ranging from only 1 primer set up to 4 primer sets. Only 1 specimen gave an amplicon of 1,365 bp, while others gave single to multiple positive amplicons in both continuous and discontinuous regions. This indicated that the amplicons did not arise from contiguous targets and might vary in insertion length. The results also confirmed that none of the shrimp were infected with WSSV. Sequencing of the PCR amplicons of expected sizes revealed sequence identity to extant WSSV genomes. This data will be used to screen the breeding stock for EVE that provide WSSV tolerance.

## 1. INTRODUCTION

Viral accommodation refers to the specific, adaptive shrimp and insect immune responses to viral infections that occur in individual cells and can lead to innocuous infections lasting up to a host’ s lifetime (see reviews (Flegel, 2020, 2022). In summary, the responses involve host production of viral copy DNA (vcDNA) fragments from viral RNA in both linear (lvcDNA) and circular (cvcDNA) forms via endogenous host reverse transcriptase (RT). This is followed by production of small interfering RNA (siRNA) and a cellular and systemic RNA interference (RNAi) responses, together with autonomous insertion of some vcDNA fragments into the host genomic DNA as endogenous viral elements (EVE). If the EVE produce antisense RNA, they may also induce a protective RNAi response. EVE in germline cells are heritable (Utari et al., 2017; Taengchaiyaphum et al., 2019).

It has recently been revealed that shrimp EVE may also give rise to cvcDNA that can be selectively isolated and sequenced as a means of screening for EVE in shrimp breeding stocks (Taengchaiyaphum et al., 2021). Based on data from whole genome sequencing of the giant tiger shrimp *Penaeus monodon*, it has been shown that EVE occur as fragments together with host transposable element sequences in host shrimp piRNA-gene-like clusters bracketed by linear repeats (Huerlimann et al., 2022; Taengchaiyaphum et al., 2022). The viral genome fragments in these clusters are scrambled in terms of reading direction and original genome position and may also redundantly overlap when mapped to the target virus genome. Thus, short read sequences from routine high throughput sequencing and assembly processes may give rise to an erroneous apparently continuous, complete or almost complete whole viral genome sequence when mapped against a database of known viral sequences.

Since white spot syndrome virus (WSSV), the causative agent of white spot disease is among the most threatening of viruses to shrimp cultivation, this study focused on cvcDNA isolation and sequencing to identify WSSV-EVE from a domesticated breeding stock of the whiteleg shrimp *Penaeus vannamei*. This stock had originated from populations exposed to WSSV for over 10 years under normal farming conditions and were used to develop a specific pathogen free (SPF) and specific pathogen tolerant (SPT) population. In the case of WSSV, selection of individual survivors of over 30g from WSSV disease outbreaks in farmed shrimp were cold challenged and only nested PCR negative animals were selected. This was followed by a long-term breeding selection process for high survival upon challenge with WSSV maintaining the SPF status under pathogen exclusion conditions. Based on the viral accommodation mechanism, we hypothesized that this stock would contain individuals with a higher prevalence of WSSV-protective EVE than might be found in randomly selected wild shrimp. Our long-term goal is to identify highly protective EVE and to develop chimeric shrimp-virus PCR primers that can be used to maintain them and possibly combine them for even higher WSSV tolerance during continued selective breeding for other commercially valuable traits.

Here we describe the first steps in working towards these goals. We reveal that cvcDNA of WSSV can be extracted from a domesticated shrimp breeding stock selected for tolerance to WSSV, and that it can be sequenced for use in designing primers for PCR detection of WSSV-EVE in individual shrimp. The information gained will facilitate screening for WSSV-EVE by PCR and testing them for protection against white spot disease (WSD) singly or in combination.

## 2. MATERIALS AND METHODS

### 2.1. Source of shrimp

DNA samples (n=50) in 95% ethanol from a whiteleg shrimp *P. vannamei* breeding stock were obtained from the National Aquaculture Group (NAQUA), Kingdom of Saudi Arabia. This shrimp breeding stock in a biosecure breeding center that was free of WSSV and has remained free of WSSV to this date (10 years period). The stock was derived from survivors of WSD outbreaks that were screened for freedom from infectious WSSV (Alday-Sanz et al., 2020) and constitutes an specific pathogen free (SPF) stock. The DNA extraction process was performed using Exgene^TM^ cell SV mini DNA extraction kits (GeneAll, Korea) and the DNA concentration was determined by Nanodrop spectrophotometer (ThermoFisher Scientific, USA). The individual DNA stock was diluted to 50ng/µl for PCR analyses and a 2 µg aliquot from each specimen was used for the circular DNA (cirDNA) extraction step.

### 2.2. Preliminary PCR screening

A preliminary PCR screening was carried out individually with a portion of all 50 DNA extracts using primers designed to detect the presence of WSSV-EVE366 as previously described (Utari et al., 2017). EVE366 was previously reported as the dominant EVE (84.8%) in the black tiger shrimp, *P*. *monodon* breeding stock of the Shrimp Genetic Improvement Center in Thailand (Utari et al., 2017). We hypothesized that EVE366 might provide some selective advantage. DNA extracts from the 50 shrimp in this study were screened for the presence of EVE366 before the DNA was pooled and subjected cirDNA isolation and sequencing.

Briefly, The PCR reaction was prepared in 12. 5 µl consisting of 1x One*Taq*^®^ PCR master mix (NEB, USA), 0. 5 µM of each forward primer (5’ ATGGGGAAGATCCTTAGAGA 3’) and reverse primer (5’CTCTTCTGGTTGCAATAG TT 3’) and 100 ng of DNA extract. The PCR amplification protocol was denaturation at 94°C for 5 min, followed by 35 cycles of 94°C, 55°C and 72°C for 30 sec each and a final extension at 72°C for 5 min. The positive PCR amplicon (122 bp) was determined by 1.5% agarose gel electrophoresis and ethidium bromide staining. To confirm the presence of circular DNA in the shrimp DNA extracts, primers specific for shrimp mitochondrial DNA were used (Pv-mt_171F: 5’ CCG TAG ACA ACG CTA CAC TAAC3’, Pv-mt_171R: 5’ATG TGA AGT AAG GGT GGA AAGG 3’).

### 2.3. Preparation of circular DNA for high throughput sequencing

Altogether 10 of the 28 DNA samples that tested positive for EVE366 were selected and pooled for cvcDNA extraction and its preparation for DNA sequencing as previously described (Taengchaiyaphum et al., 2021) (**Fig. 1**). Briefly, two micrograms of the pooled shrimp genomic DNA was aliquoted and subjected to linear DNA digestion using plasmid-safe DNase (Epicentre, UK) for 4 days. The remaining cirDNA was precipitated by ethanol-acetate and then the concentration was determined by Qubit Fluorometer (Invitrogen, USA).

**Figure 1.**
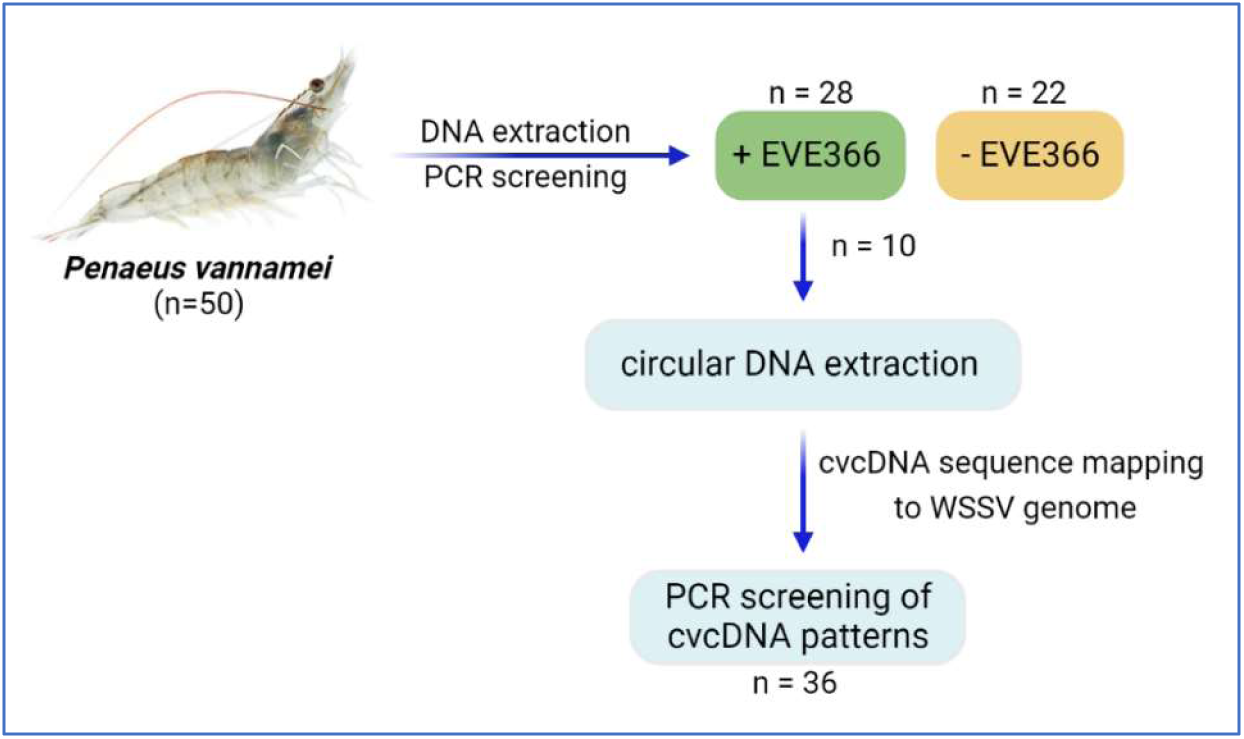
Diagram showing the work plan for isolation, sequencing and analysis of samples of putative cvcDNA derived from the 50 samples of shrimp broodstock.

### 2.4. Sequencing and analysis of pooled cvcDNA

A portion of the total cvcDNA preparation was subjected to rolling circle amplification (RCA) by using a Repli-G mini kit (Qiagen, Germany) following the manufacturer’s protocol. The RCA product was precipitated and DNA concentration determined by Qubit Fluorometer before the product was evaluated by 1. 5% agarose gel electrophoresis prior to being send for high throughput sequencing. Sequencing of the RCA product was performed and analyzed by Novogene (Novogene Corporation Inc., Singapore). In brief, twenty micrograms of RCA product were randomly fragmented by sonication, then DNA fragments were end polished, A-tailed, and ligated with the full-length adapters of Illumina sequencing. This was followed by further PCR amplification with P5 and indexed P7 oligos. The PCR products as the final construction of the libraries were purified with the AMPure XP system. Then libraries were checked for size distribution by Agilent 2100 Bioanalyzer (Agilent Technologies, CA, USA), and quantified by real-time PCR (to meet the criteria of 3 nM). For the data analysis step, raw data filtering of low quality reads was carried out to generate clean data. SPAdes de novo assembly software (version 3.11.1) was used to assemble the sample reads from the clean Data against a WSSV reference genome (GenBank accession no. AF440570). This provided a library of WSSV sequences derived from putative WSSV-cvcDNA fragments that could be mapped against the reference genome and allow the design of primers to screen the original, individual DNA extracts for the presence of homologous EVE/cvcDNA.

### 2.5. Detection of cvcDNA-related fragments in the shrimp genomic DNA extract

A total of 36 from 50 original shrimp DNA samples with sufficient original DNA remaining (section 2.1) were subjected to PCR using primers designed from the WSSV cvcDNA sequencing process to screen for homologous WSSV-EVE. The specific primers were designed based on high-frequency-read fragments (HFRF) determined by mapping raw WSSV reads against the reference genome as described in the previous section. PCR reactions were carried out using One*Taq* PCR mastermix and the amplification conditions were similar to those previously described in section 2.2. The list of primers used in this step is shown in **Table 1**.

**Table 1.**
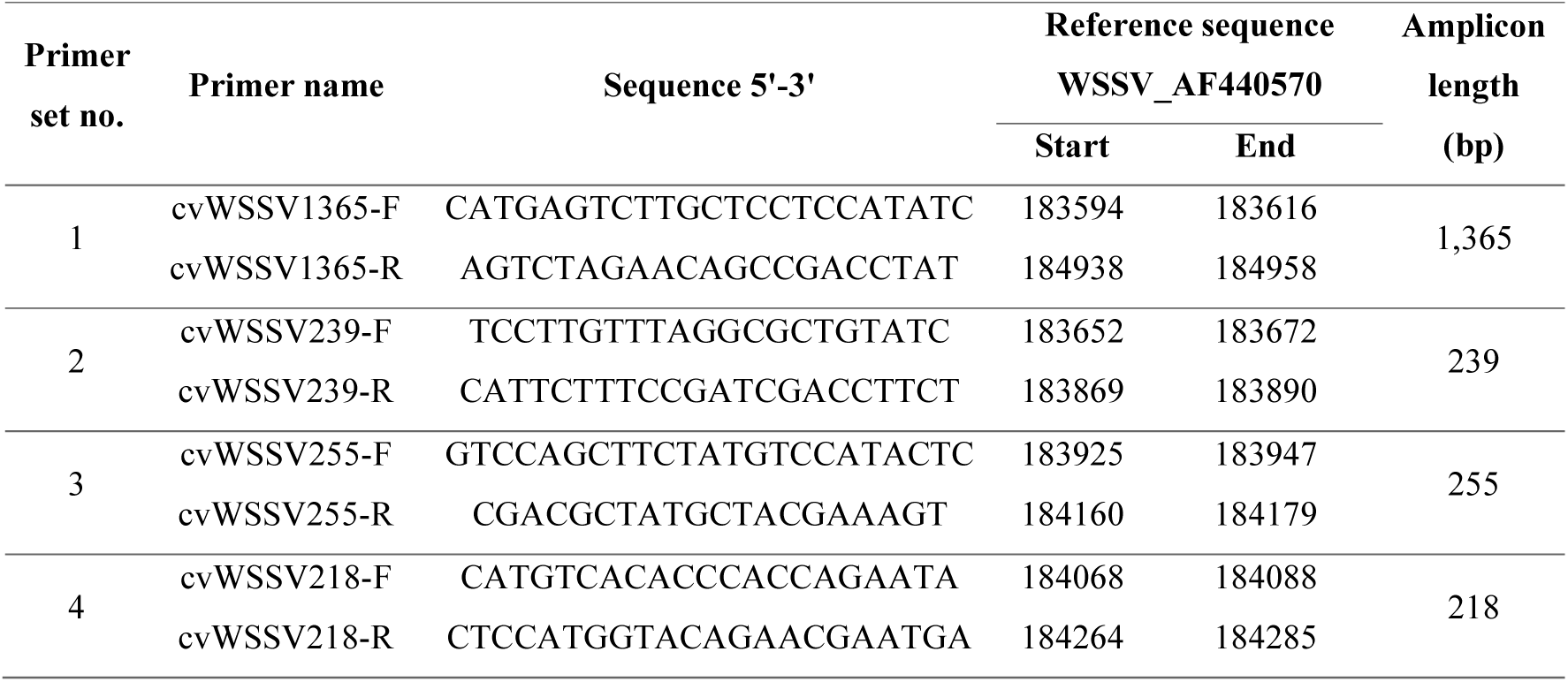
Primer sets designed to test for the presence of HFRF fragments of WSSV-EVE in individual shrimp from the NAQUA stock. These are shown diagrammatically in **Fig. 5**. Note that the amplicon lengths are those predicted using the genome of AF440570 as the template. Amplicon length may differ with DNA templates of individual shrimp specimens. Some of the primers are used in more than one primer set. The primer sets are numbered from 1 to 4 based on the sequential order of the forward primers in the GenBank reference sequence AF440570.

### 2.6. PCR amplicon sequencing

The PCR amplicons were purified, then ligated to pGEM-T easy vector (Promega, USA) prior to transforming *Escherichia coli*-DH5α (ThermoFisher Scientific, USA). Three cloned plasmids for each amplicon were sent for DNA sequencing (ATGC Co., Ltd., Thailand). Sequence comparisons were achieved using Clustal Omega (https://www.ebi.ac.uk/Tools/msa/clustalo/) using the reference WSSV genome GenBank accession no. AF440570.

## 3. RESULTS

### 3.1. Preliminary screening for EVE366 in NAQUA broodstock samples

A set of 50 DNA samples from the *P. vannamei* NAQUA breeding stock proven to be free of WSSV were tested for WSSV-EVE_366_ previously found to be the most prevalent of 4 EVE known to occur in *P. monodon* from Thailand (Utari et al., 2017). The WSSV-EVE366 forward primer is located at positions 251,608 to 251,635 in GenBank record AF440570. This was a preliminary test based on the assumption that PCR-positive DNA samples might possibly contain WSSV-cvcDNA as indicators for the presence of WSSV-EVE366. As a first step, all 50 of the DNA extracts were subjected to PCR tests for the presence of mitochondrial DNA as a positive control for the presence of intact circular DNA in the extracts (**Fig. 2A**).

**Figure 2.**
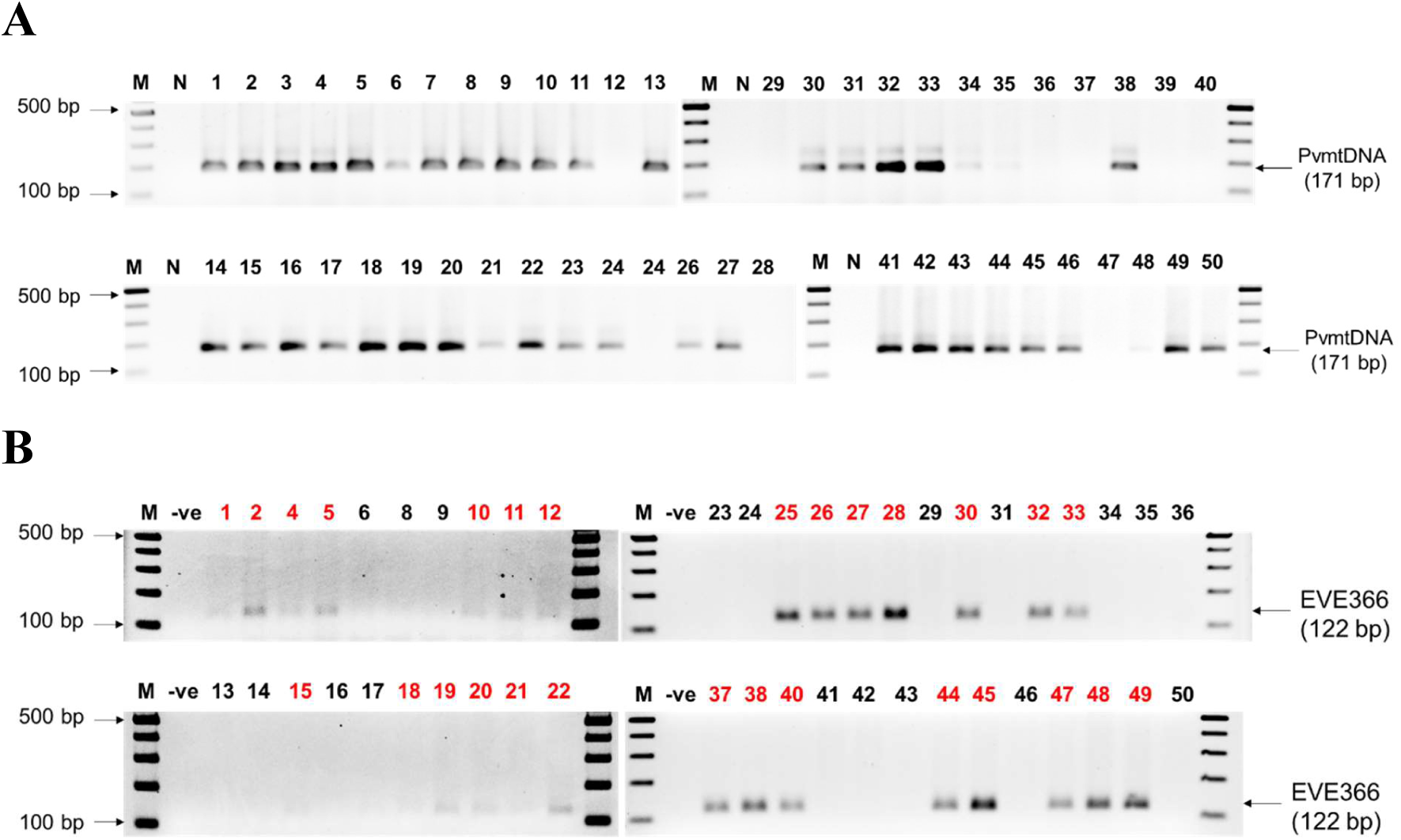
Preliminary PCR tests with the 50 DNA extracts. **A.** Inverted agarose gel photomicrographs showing positive PCR test results for the presence of circular shrimp mitochondrial DNA in all 50 samples. **B.** Inverted photomicrographs of PCR amplicon bands indicating the presence of the WSSV-EVE366 target sequence (red numbers) for 28 of the 50 DNA samples.

When the same samples were also subjected to PCR testing for the presence of WSSV-EVE366 28 of the 50 DNA samples gave positive results (**Fig. 2B**). These results were sufficiently encouraging to go ahead with cvcDNA extraction followed by sequencing for all samples in lots of 10 samples each. The results for the WSSV-EVE_366_ PCR tests for all the samples are shown in tabular form in **Table 2**.

**Table 2.** Summary table showing identities and characteristics of the 50 samples tested for mtDNA and for EVE366. The numbers in the column labeled “ Pool DNA set” indicate the samples pooled in sets for cvcDNA extraction followed by sequencing. In this publication, we present data from DNA Pool 1 only (gray shade indicates the samples included in the pool).

| Sample | mtDNA | EVE366 | Pool DNA set | Sample | mtDNA | EVE366 | Pool DNA set |
| --- | --- | --- | --- | --- | --- | --- | --- |
| 1 | + | + | 1 | 26 | + | + | 2 |
| 2 | + | + | 1 | 27 | + | + | 2 |
| 3 | + | - | 3 | 28 | + | + | 2 |
| 4 | + | + | 1 | 29 | + | + | 4 |
| 5 | + | + | 1 | 30 | + | - | 2 |
| 6 | + | - | 3 | 31 | + | + | 4 |
| 7 | + | - | 3 | 32 | + | - | 2 |
| 8 | + | - | 3 | 33 | + | + | 2 |
| 9 | + | - | 3 | 34 | + | + | 4 |
| 10 | + | + | 1 | 35 | + | - | 4 |
| 11 | + | + | 1 | 36 | + | - | 4 |
| 12 | + | + | 1 | 37 | - | - | 2 |
| 13 | + | - | 3 | 38 | - | + | 2 |
| 14 | + | - | 3 | 39 | + | + | 4 |
| 15 | + | + | 1 | 40 | - | - | 2 |
| 16 | + | - | 3 | 41 | - | + | 3 |
| 17 | + | - | 3 | 42 | + | - | 3 |
| 18 | + | + | 1 | 43 | + | - | 3 |
| 19 | + | + | 1 | 44 | + | - | 4 |
| 20 | + | + | 2 | 45 | + | + | 4 |
| 21 | + | + | 2 | 46 | + | + | 3 |
| 22 | + | + | 2 | 47 | + | - | 4 |
| 23 | + | - | 3 | 48 | + | + | 4 |
| 24 | + | - | 4 | 49 | + | + | 4 |
| 25 | + | + | 2 | 50 | + | + | 3 |

The reasons for differences in the intensity of the target amplicon of 239 bp from specimen to specimen are unknown. Since total template DNA was equal for all specimens, it is possible that darker intensity in some bands may indicate the presence of more than one copy of the target sequence in the genome of the source specimen or possible amplification of that EVE in the form of cvcDNA (Taengchaiyaphum et al., 2021). For example, up to 3 copies of the target sequence of a PCR detection method for the shrimp virus IHHNV were found in an EVE cluster on chromosome 7 of a whole genome record for the giant tiger shrimp *Penaeus monodon* (Taengchaiyaphum et al., 2022). As has been shown with IHHNV-EVE, the same region of virus genome may be observed in both sense and anti-sense directions.

The results obtained for the presence of mitochondrial DNA (host circular DNA representative) and for the presence of EVE366 in all 28 samples encouraged us to continue with extraction of cvcDNA to be pooled and sent for NGS analysis. For the first pool of DNA extracts, we selected the first 10 extracts (**Table 2** gray outline) that gave positive PCR test results for WSSV-EVE366.

### 3.2. WSSV-EVE confirmed by sequencing of cvcDNA pool 1

When short-read sequences from the samples were aligned with the whole WSSV genome sequence of an extant type of lethal WSSV (GenBank record AF440570), those with high sequence homology to WSSV (99 to 100%) mapped to positions located across the whole WSSV genome of 307, 287 base pairs (bp) (**Table 3 & Fig. 3**).

**Figure 3.**
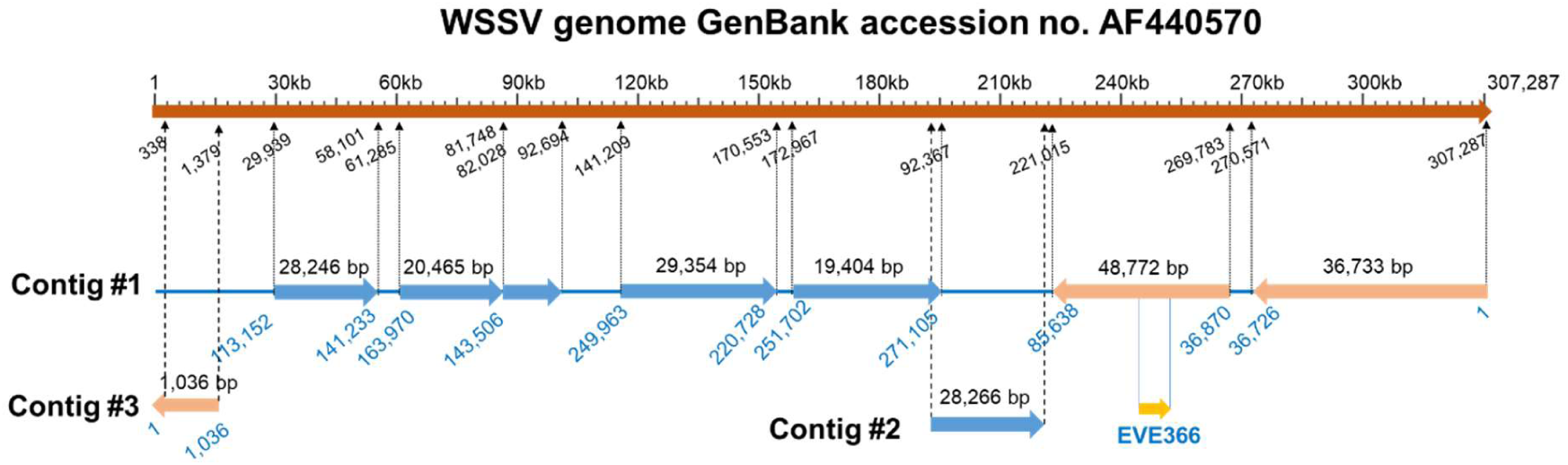
Map of the location of various contigs constructed from the NGS data and showing distribution over the whole WSSV genome for nucleic acid sequences with high homology (up to 99% identity to the reference GenBank record) to an existing, lethal type of WSSV (GenBank record AF440570). Altogether, these putative EVE sequences accounted for approximately 69% of the whole WSSV genome.

**Table 3.**
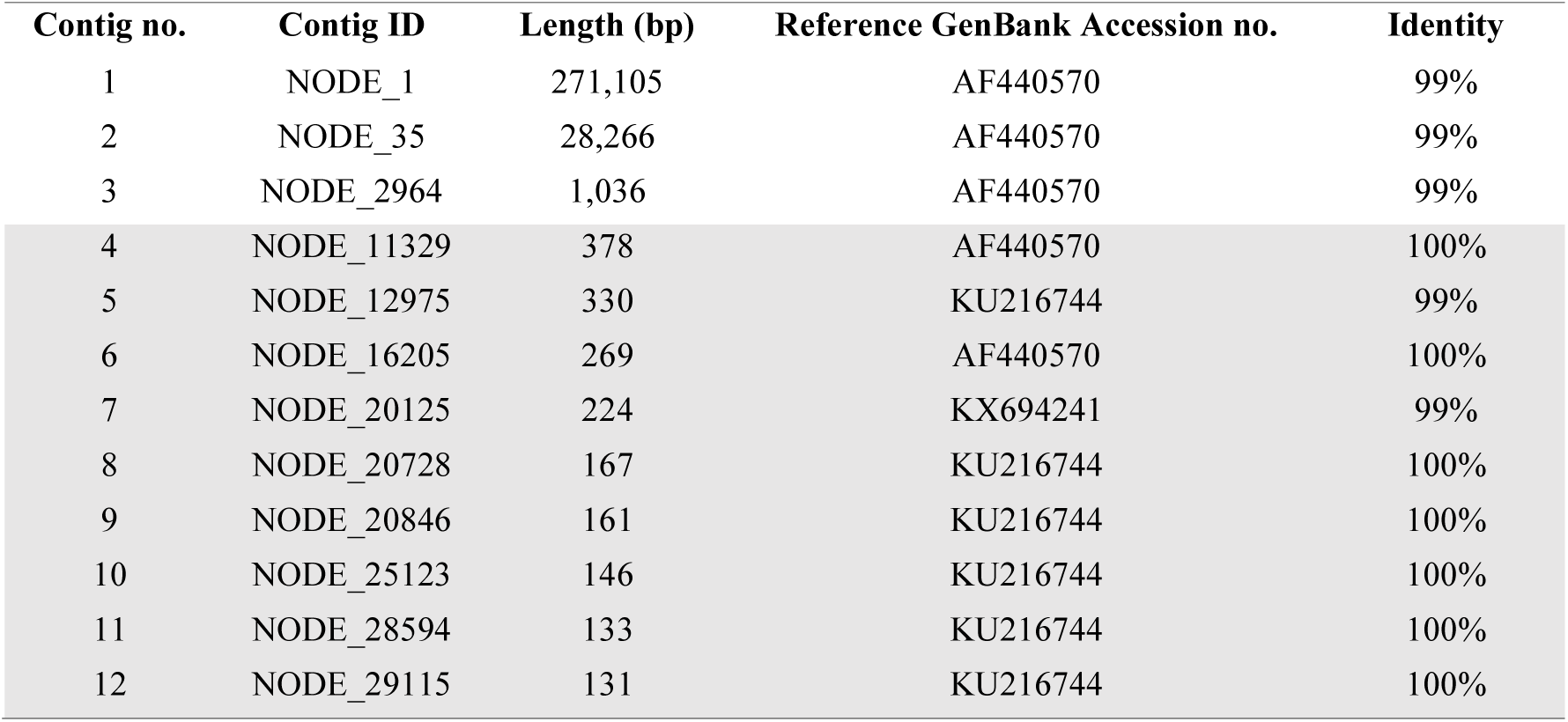
List of sequences assembled from pooled cvcDNA sample set 1. The contigs no.4-12 (outlined in gray) matched repetitive sequences found in the WSSV genome reference. Mapping of contigs no.1-3 to the WSSV genome database is shown in Fig. 3.

| Contig no. | Contig ID | Length (bp) | Reference GenBank Accession no. | Identity |
| --- | --- | --- | --- | --- |
| 1 | NODE_1 | 271,105 | AF440570 | 99% |
| 2 | NODE_35 | 28,266 | AF440570 | 99% |
| 3 | NODE_2964 | 1,036 | AF440570 | 99% |
| 4 | NODE_11329 | 378 | AF440570 | 100% |
| 5 | NODE_12975 | 330 | KU216744 | 99% |
| 6 | NODE_16205 | 269 | AF440570 | 100% |
| 7 | NODE_20125 | 224 | KX694241 | 99% |
| 8 | NODE_20728 | 167 | KU216744 | 100% |
| 9 | NODE_20846 | 161 | KU216744 | 100% |
| 10 | NODE_25123 | 146 | KU216744 | 100% |
| 11 | NODE_28594 | 133 | KU216744 | 100% |
| 12 | NODE_29115 | 131 | KU216744 | 100% |

We now know that the sequences of the WSSV contigs are very likely an artificial result of the sequence assembly process that uses overlapping small-read sequences assembled against a reference sequence, rather than an actual contiguous sequence in the target genome. This was clearly revealed for EVE of the shrimp virus IHHNV in full shrimp genome sequences developed using long-read technology (Huerlimann et al., 2022; Taengchaiyaphum et al., 2022). The EVE fragments of the IHHNV genome occurred in clusters scrambled (in location and reading direction), some with overlapping sequences that resulted in a cumulative sum of IHHNV-EVE sequence lengths that exceeded the whole IHHNV genome length.

### 3.3. High frequency read fragments mapped to a minute part of the WSSV genome

Most of the cvcDNA fragments with high homology to WSSV occurred at low prevalence, while high frequency read fragments (HFRF) mapped to only a small part of the genome covering approximately 1,400 bp (**Fig. 4A and B**). Since the source breeding stock was developed from uninfected survivors of WSSV disease outbreaks, the data suggested that the high frequency EVE sequences from that region of the WSSV genome may have been naturally selected in initial outbreak survivors and then further in challenge tests designed to select the shrimp most tolerant to WSSV challenge.

**Figure 4.**
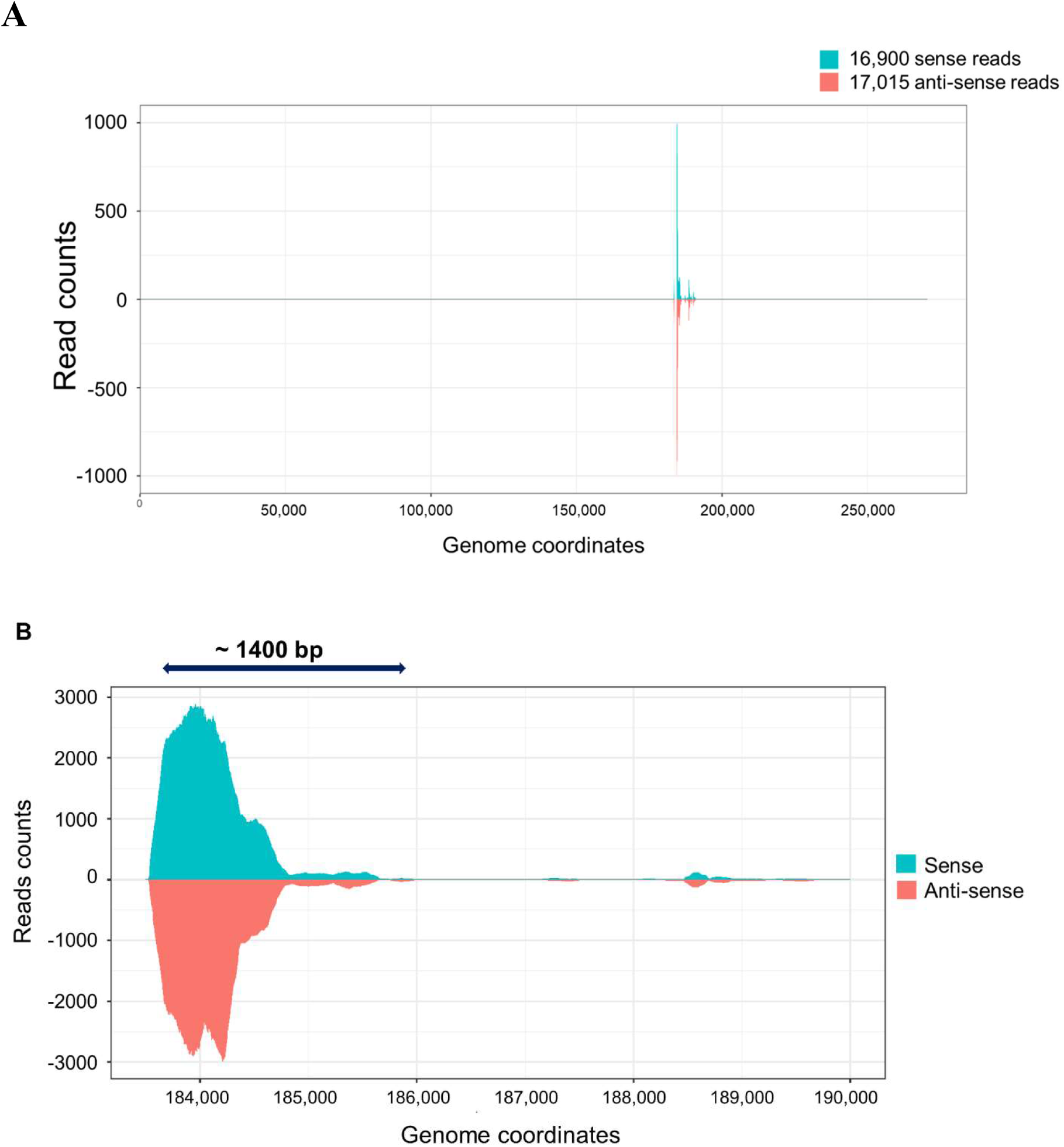
Various contigs constructed from the NGS data and showing DNA fragments with high homology to the genome sequence of an existing lethal type of WSSV (GenBank record AF440570) scattered over the whole WSSV sequence. (A) Whole genome overview showing reads from 1 upwards. (B) Enlarged map focused on the map region of approximately 1,400 bp showing high frequency read fragments (HFRF).

While looking at Fig. 4, it is important to keep in mind that low frequency hits were found distributed over the whole WSSV genome (Fig. 3). In addition, Fig. 4 gives no indication as to how the fragments used to prepare this map were distributed in the genomes of the 10 individual shrimp used to prepare the DNA pool.

### 3.3. High individual shrimp EVE variation revealed by HFRF-PCR analysis

Based on the hypothesis that high-frequency-read fragments (HFRF) indicate positive selection due to induced WSSV tolerance in carriers, PCR primer sets were designed (**Table 1**) to cover the 1,400 bp region shown in **Fig. 4**. These are shown diagrammatically compared to the GenBank reference sequence AF440570 in **Fig. 5**.

**Figure 5.**
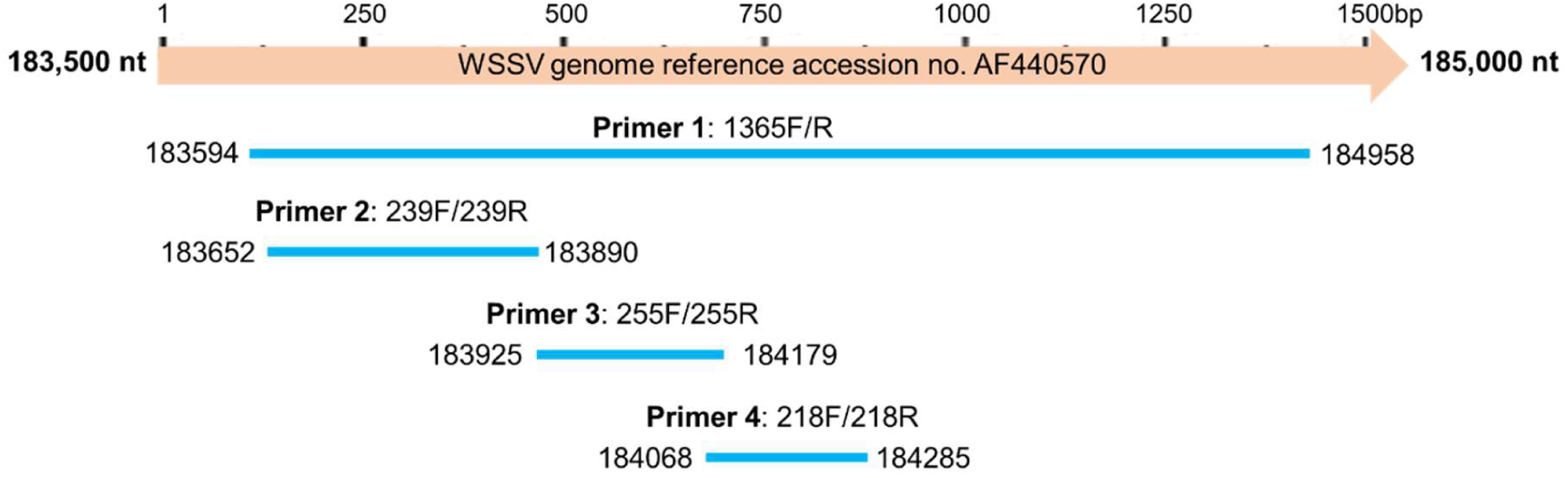
Diagram of the amplicons that would be expected using the primer sets shown in Table 1.

In Fig. 4, the genome of GenBank record AF440570 was used as the template to design the primers illustrated in Fig. 5. In reality, the amplicons obtained using genomic DNA from NAQUA shrimp samples might differ. Thus, all such amplicons would have to be sequenced individually. Also note that Primer sets 2 to 4 all fit within the map range covered by Primer set 1. Thus, if an amplicon obtained using Primer set 1 with a shrimp sample arose from a single, continuous EVE that matched exactly with the sequence of AF440570, it would include the sequences of all the remaining Primer sets and would thus give positive PCR test results for all 4 primer sets. On the other hand, a positive PCR test result for Primer set 2 alone, for example, would not indicate the presence of adjacent sequences related to the sequences targeted by other primer sets.

Of the 50 shrimp samples shown in Table 2 above, only 36 samples had sufficient DNA remaining to carry out PCR reactions with all 4 primer sets. When this was done with all 36 samples (144 reactions), the results shown in **Table 4** were obtained and are summarized pictorially in **Fig. 6**. From Table 4, it can be seen that only 4 of the 36 specimens gave negative test results using the 4 PCR methods, and that 32 out of the 36 shrimp (89%) gave positive PCR test results with at least one of the WSSV sequence-fragments targeted by the 4 primer sets. This pattern is similar to that obtained when overlapping primers were used to screen 40 shrimp that had been selected using a WOAH-recommended detection method for the virus IHHNV (Saksmerprome et al., 2011). Only 3 of the 40 shrimp specimens (7%) were positive for IHHNV using all the overlapping primers and confirming IHHNV infections. The other 37 (93%) showed only fragments of the IHHNV genome and had thus given false-positive test results for IHHNV using the single, recommended method.

**Figure 6.**
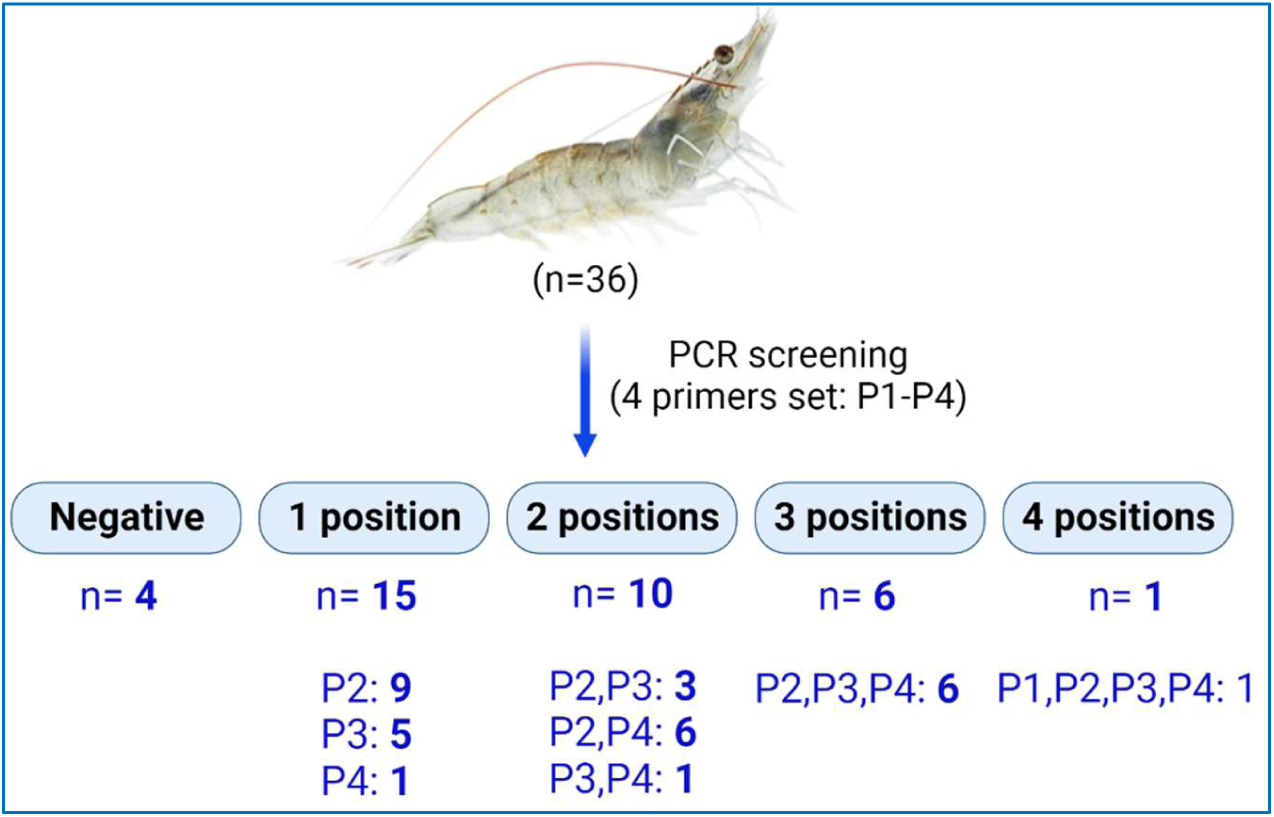
Pictorial results of PCR testing using 4 primer sets with 36 specimens from the breeding stock.

**Table 4.**
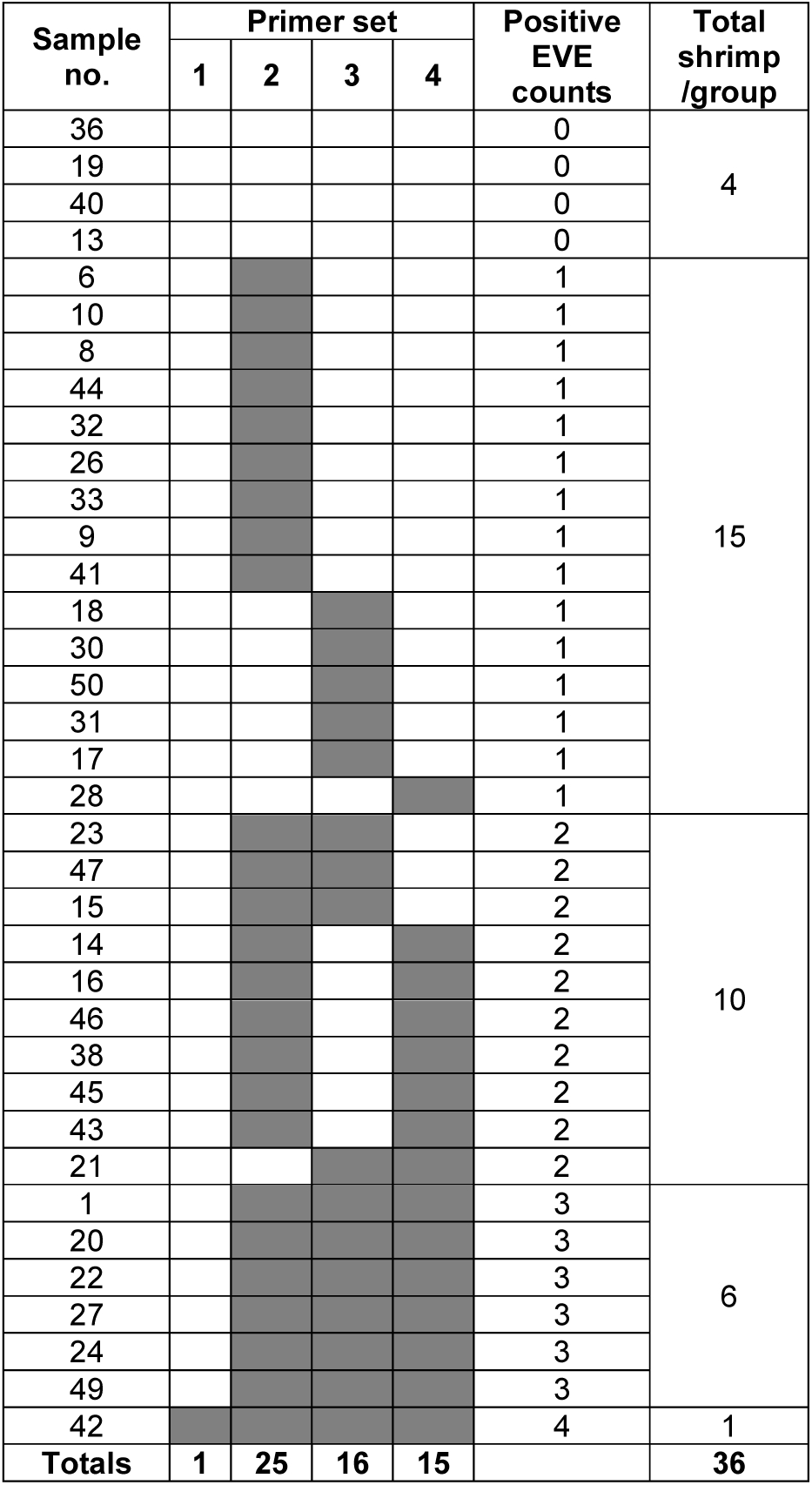
Results of PCR testing using 4 primer sets with 36 specimens from the breeding stock. The gray rectangles indicate positive PCR test results. The table is arranged so that shrimp with only one positive test result is at the top followed by those with increasing numbers of positive test results. Total positives for each specimen (0 to 4) are indicated on the right side of the table together with the number of specimens giving the same number of positive PCR tests (1-4). The total number of positive results for each primer set are given at the bottom of the table.

Only one of the 36 shrimp specimens (#42) carried an EVE with a sequence corresponding to the whole 1,365 bp high frequency region mapped to the WSSV genome in **Fig. 4**. In addition, except for specimen #42 that was positive with all 4 primer sets, the failure of the remaining 35 specimens to give positive results with 1 to 3 of the 4 primers sets provided additional confirmation that those specimens were not infected with WSSV. Sequencing of the amplicon from specimen #42 and sequencing a random selection of the amplicons from the other primer sets 2, 3 and 4 revealed that all had very high sequence identity (99 to 100 %) to an extant type of WSSV (Supplementary file 1).

Of particular interest were the HFRF primer sets that gave positive PCR test results with a high proportion of the 36 tested shrimp (bottom line in Table 4). These were primer sets 2 (25/36 = 69.4%) and 3 (16/36 = 44%). These will constitute the preliminary test primers used to screen the breeding stock to obtain individuals suitable to arrange crosses for testing the protective potential of these 2 target EVE.

### 3.4. Multiple PCR amplicon bands were obtained with many of the 36 specimens

Many of the shrimp DNA samples gave multiple PCR amplicons with the 4 primer sets described in Section 3.4. An example PCR gel obtained using DNA templates for 12 of the 36 specimens with Primer Set 2 (i.e., the set with 69% of the 36 samples positive) is shown in **Fig. 7**. The intensity of the amplicon bands for the expected target sequence of 239 bp varied from sample to sample (e. g., 1,8,10, 13, 14, 15, 16, 20). At the same time, some of these 239 bp-positive samples (e. g.,1, 13), also showed an additional amplicon band or bands of lower intensity and longer lengths (asterisks).

**Figure 7.**
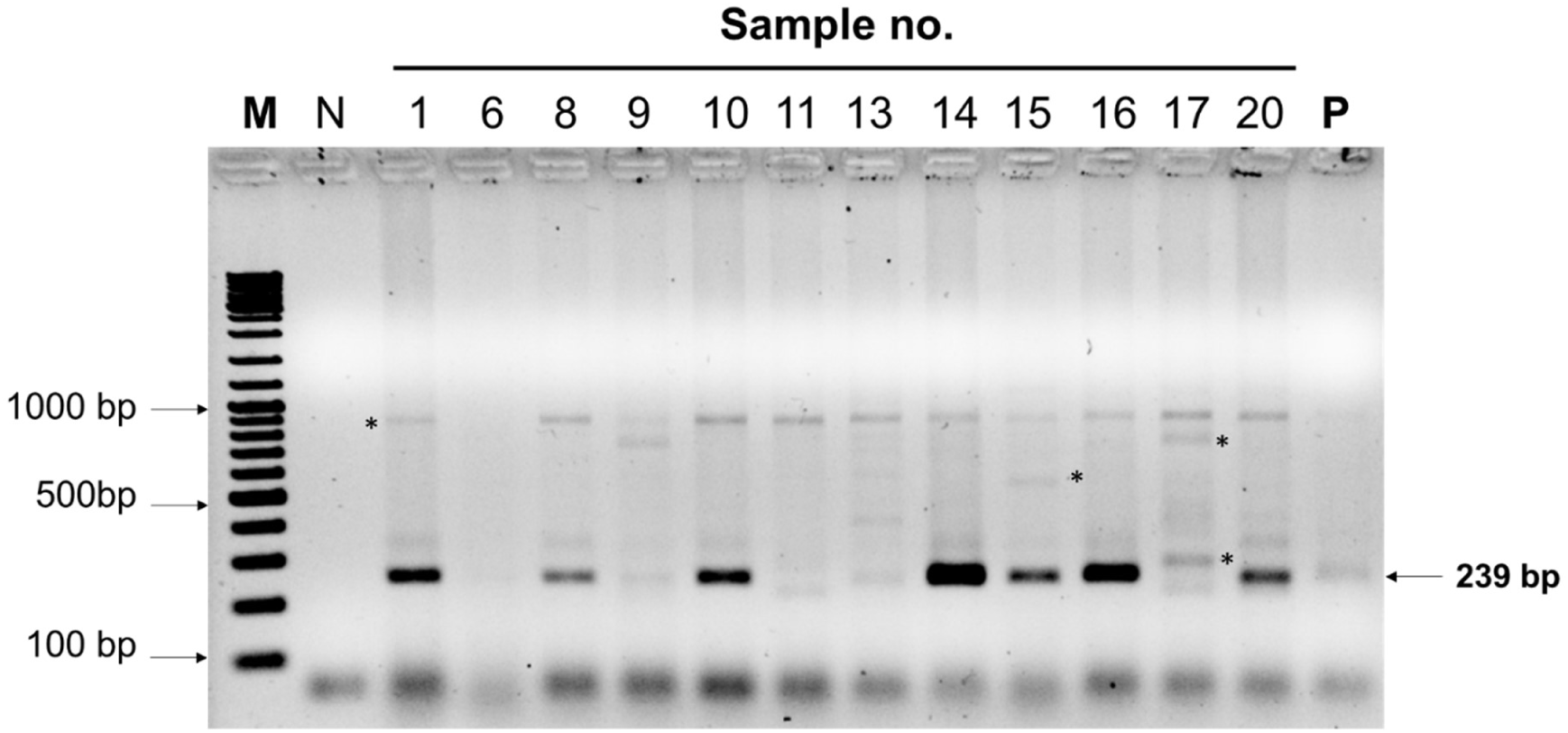
Inverted photograph of an example agarose gel for 12 out of 36 shrimp DNA specimens tested with Primer Set 2 (positive results for 69% of the 36 specimens). The expected amplicon of 239 bases was obtained with variable band intensity from specimen to specimen. In addition, many of these specimens also gave one or more additional but fainter bands of longer length (asterisks), while other specimens that did not give a band for 239 bases, did give similar bands for longer amplicons.

The reasons for differences in the intensity of the target amplicon of 239 bp from specimen to specimen are unknown. Since total template DNA was equal for all specimens, it is possible that darker intensity in some bands may indicate the presence of more than one copy of the target sequence in the genome of the source specimen or possible amplification of that EVE in the form of cvcDNA (Taengchaiyaphum et al., 2021). For example, up to 3 copies of the target sequence of a PCR detection method for the shrimp virus IHHNV were found in an EVE cluster on chromosome 7 of a whole genome record for the giant tiger shrimp *Penaeus monodon* (Taengchaiyaphum et al., 2022). As shown in IHHNV inserts, the same region of virus genome can be observed by both sense and anti-sense directions.

Testing of the possibility of bi-direction and multiple insertion of virus sequences was performed by using single primers with the same DNA extracts as used with the normal 3’ and 5’ primer pairs. Obtaining amplicons with these tests would indicate the presence of palindromic sequences. Indeed, the PCR result showed that some samples gave PCR amplicons using only one primer (**Fig. 8**). This result may be evidence for multiple insertions of inverted virus fragments in the shrimp genome.

**Figure 8.**
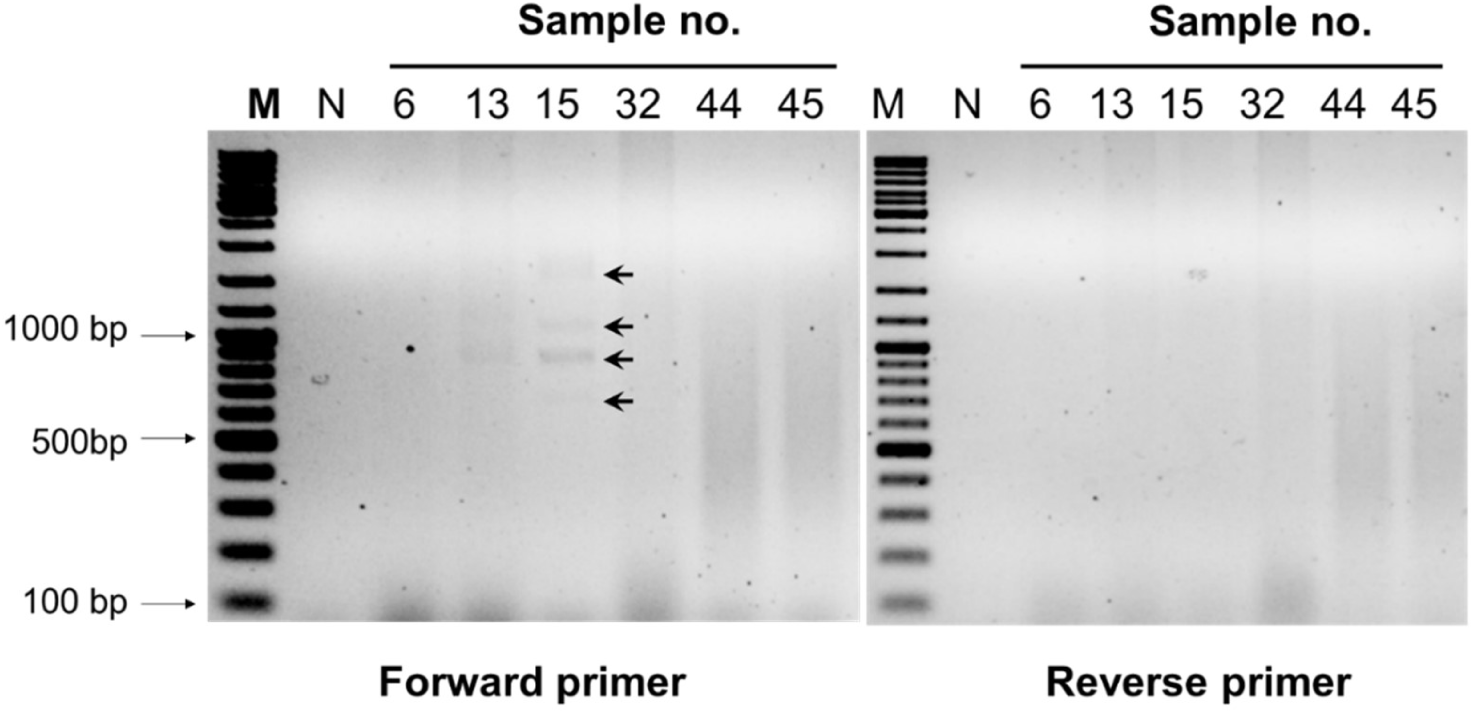
Example of amplicon bands obtained in PCR reactions using only one or the other primer from Primer set 2 with DNA extracts from 6 of our sample shrimp. Unexpected PCR amplicons (arrows) can be observed with sample 15.

### 3.5. Confirmation of WSSV-EVE by PCR and amplicon sequencing

Selected PCR amplicons were collected from individual samples to confirm nucleotide sequences. The PCR amplicons were purified, then ligated to pGEM-T easy vector prior to transforming *Escherichia coli*-DH5α. The recombinant plasmids were purified and then sent for DNA sequencing using T7 and SP6 primers. The consensus sequences generated were aligned with the corresponding WSSV genome regions based on GenBank reference no. AF440570 (**see Supplementary file 1**). The DNA sequences of four samples that gave positive amplicons using 4 primer sets are illustrated. The sequencing results showed that all positive amplicons had 99 to 100% sequence identity with the WSSV reference genome (**Table 5**).

**Table 5.** List of PCR amplicons from primers P1 to P4 sent for DNA sequencing and % identity of their sequences to WSSV genome sequence AF440570.

| Sample no. | Primer set no. |  |  |  |
| --- | --- | --- | --- | --- |
|  | 1<br>(1,365 bp) | 2<br>(239 bp) | 3<br>(255 bp) | 4<br>(218 bp) |
| 6 |  | 100% IDs |  |  |
| 30 |  |  | 100%IDs |  |
| 42 | 99% IDs |  |  |  |
| 49 |  | 100% IDs |  | 100% IDs |

## 4. DISCUSSION

We isolated cvcDNA from pooled DNA extracts from 10 shrimp, sequenced it and used PCR primer sets based on resulting sequences that matched a WSSV reference genome to test DNA extracts from 36 individual shrimp specimens. Variable PCR amplicons with high sequence identity to an existing type of WSSV were obtained from most of the specimens. Some of these amplicons were associated with host transposable element sequences, proving that the amplicons arose from variable numbers and types of EVE in the genomes of the individual shrimp specimens.

Our ultimate objective is to construct a library of chimeric primer sets with one primer based on the WSSV sequence of each EVE and the other on the host sequence contiguous with that EVE in the host genome. This will help in determining the genome location of each individual EVE and facilitate monitoring its presence or absence during stock breeding and selection. The chimeric amplicon or a chimeric part of it might also be used in a southern blot of genomic DNA to test for the presence of one or more than one copy in the host genome. Alternatively, this might be done using fluorescent *in situ* hybridization (FISH) with chromosome smears. This would facilitate breeding tests to determine the efficacy of each natural EVE construct in promoting tolerance to WSSV infection and would facilitate the preservation and joining of protective EVE combinations in individual broodstock specimens.

Thus, this publication simply describes the first step in the process of discovering the presence of EVE for WSSV in an SPF breeding stock of the whiteleg shrimp *P. vannamei* in preparation for selection of shrimp broodstock carrying various HFRS EVE combinations for breeding. After breeding, the offspring of the various crosses will be allowed to grow sufficiently to allow tagging and collection of pleopod samples from individual shrimp for nucleic acid extraction and storage prior to challenge with WSSV. After WSSV challenge, we will be able to assess the relationship between EVE-expression profiles and survival or death. Hopefully, this process will allow for selection of breeding stocks with improved tolerance or resistance to WSSV.

## Acknowledgements

This work was funded by the National Research Council of Thailand (NRCT): High-Potential Research Team Grant Program, grant no. N42A650869 (to Kallaya Sritunyalucksana), the National Aquaculture Group (NAQUA), Kingdom of Saudi Arabia. It was also supported by Mahidol University under the New Discovery and Frontier Research Grant 2023 (Grant no. FF-056/2566) and the NSRF via the Program Management Unit for Human Resources & Institutional Development, Research and Innovation (Grant No: B05F640137).

## Ethics statement

This work followed Thailand’s laws for ethical animal care under the Animal for Scientific 83 Purposes ACT, B.E. 2558 (A.D. 2015) under project approval number BT-Animal document no. 06/2566) from the National Center for Genetic Engineering and Biotechnology (BIOTEC), National Science and Technology Development Agency (NSTDA), Thailand.

## Authors contributions

ST, VAS, TWF and KS, research conceptualization; ST, JS, PW, SS, JM, IIU, MBC, and MMA, Sample preparation, molecular detection and data analysis; ST, VAS, TWF and KS, data analysis and interpretation and manuscript preparation

## Supplementary file_1

1. Multiple sequence alignment of DNA sequences obtained from PCR amplicon DNA sequencing of sample no. 42 based on primer set P1 amplification. The sequences no. C1-C3 represent selected positive clones. The nucleotide positions for reference sequence are between 183,594 – 184,958 nucleotides according to GenBank accession no. AF440570. 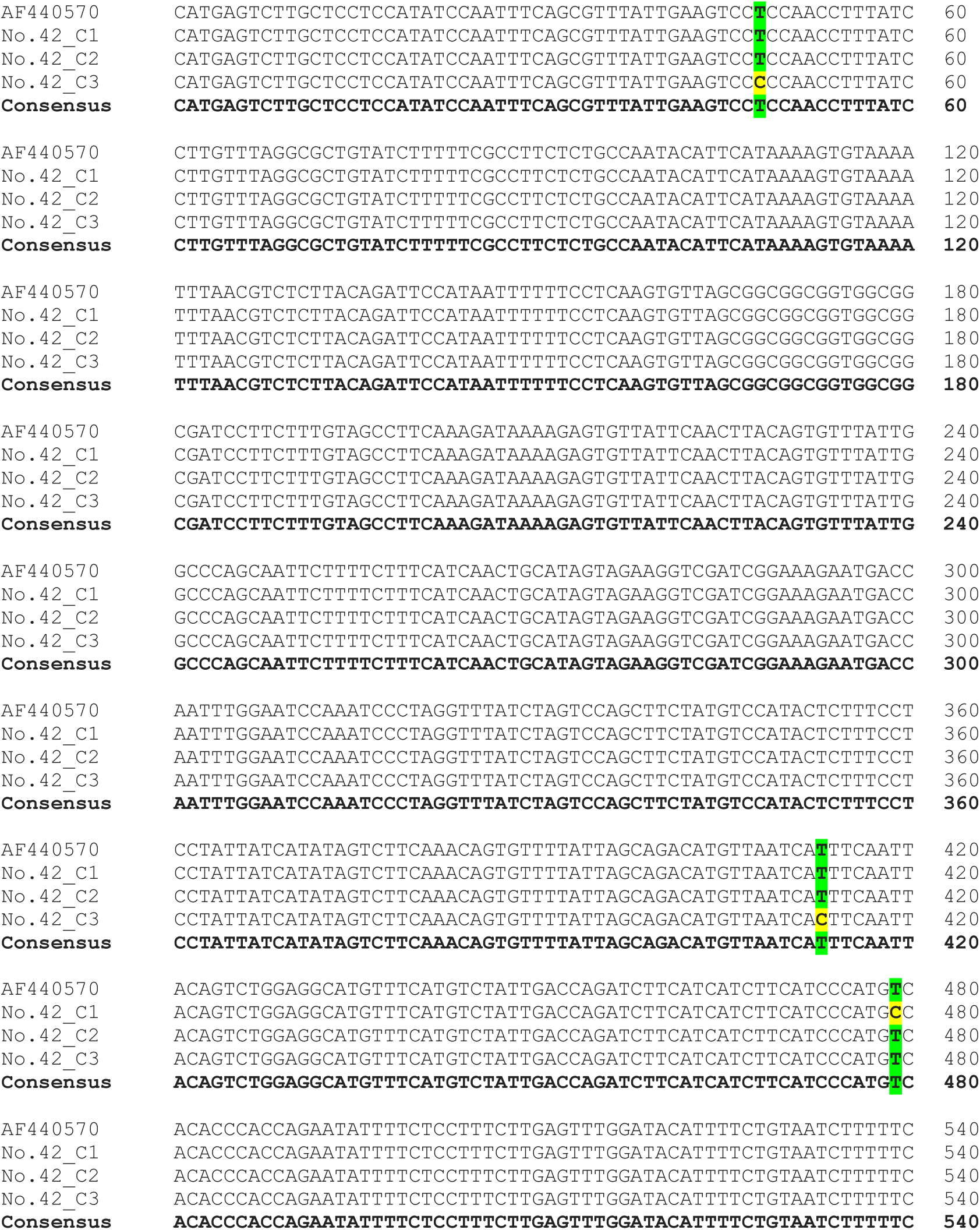
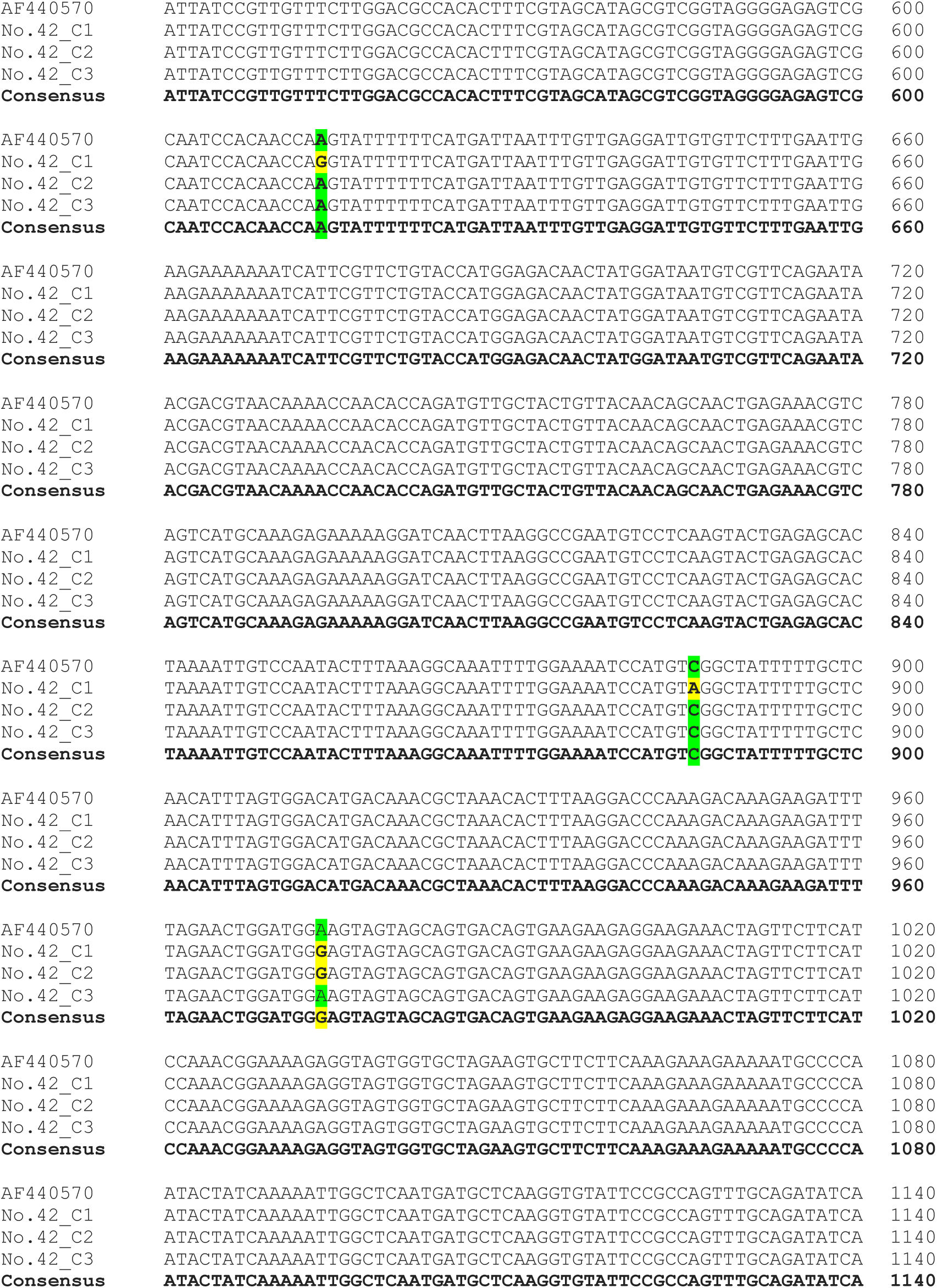
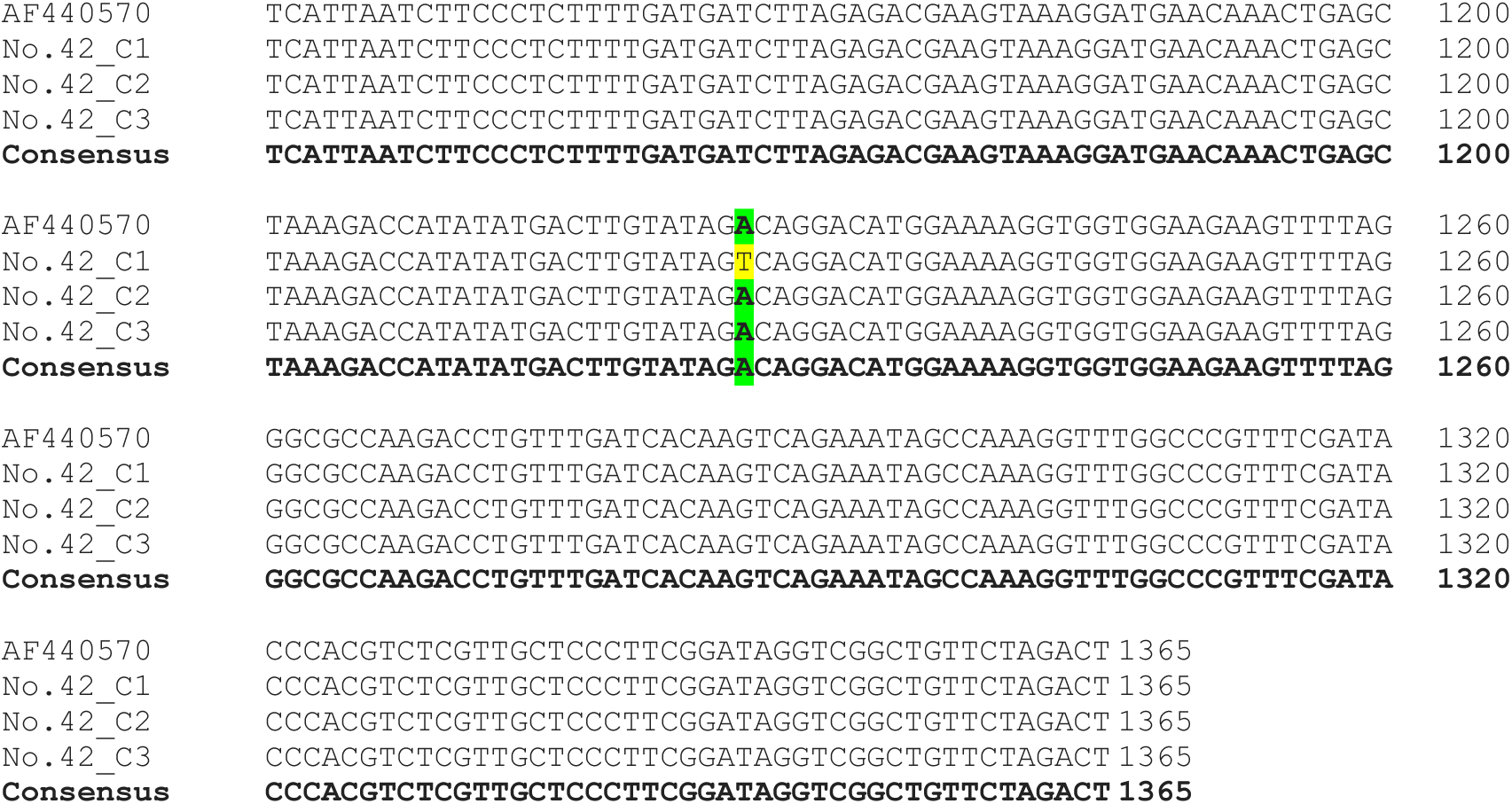
2. Multiple sequence alignment of DNA sequences obtained from PCR amplicon DNA sequencing of sample no. 6 (gray shade) and 49 based on primer set P2 amplification. The sequences no. C1-C3 represent selected positive clones. The nucleotide positions for reference sequence are between 183,652 – 183,890 nucleotides according to GenBank accession no. AF440570. 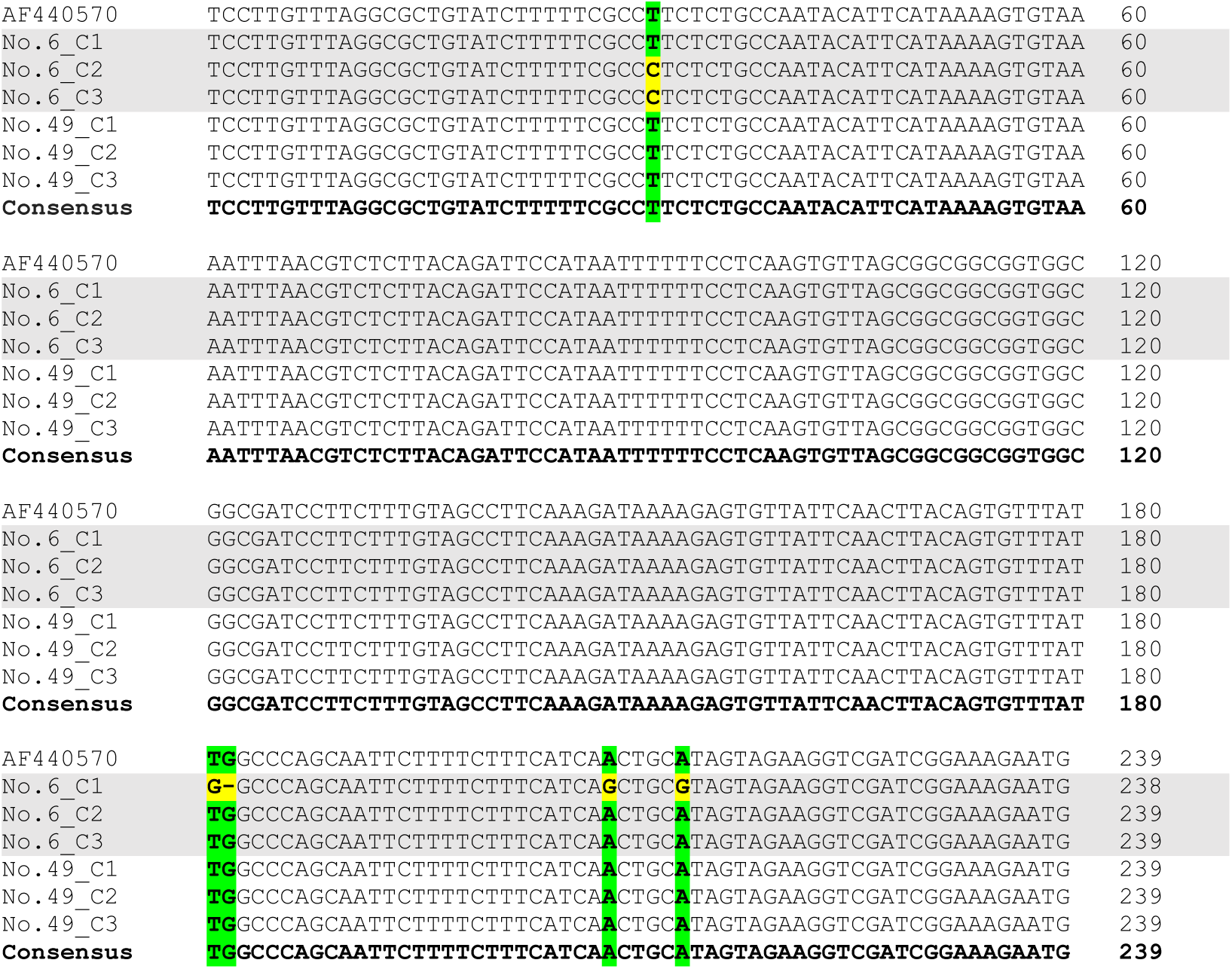
3. Multiple sequence alignment of DNA sequences obtained from PCR amplicon DNA sequencing of sample no. 30 based on primer set P3 amplification. The sequences no. C1-C3 represent selected positive clones. The nucleotide positions for reference sequence are between 183,925 – 184,179 nucleotides according to GenBank accession no. AF440570. 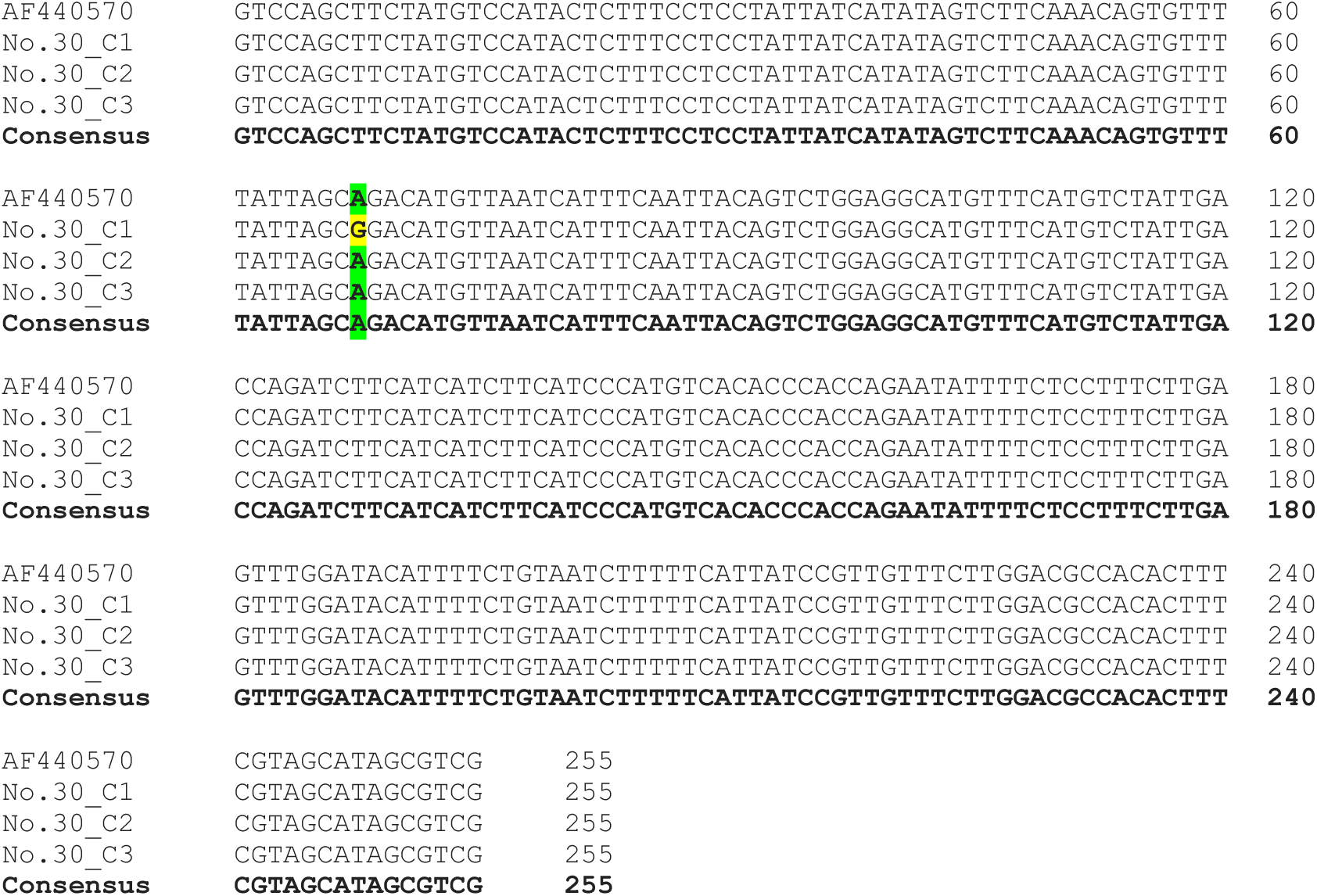
4. Multiple sequence alignment of DNA sequences obtained from PCR amplicon DNA sequencing of sample no.49 based on primer set P4 amplification. The sequences no. C1-C3 represent selected positive clones. The nucleotide positions for reference sequence are between 184,068 – 184,285 nucleotides according to GenBank accession no. AF440570. 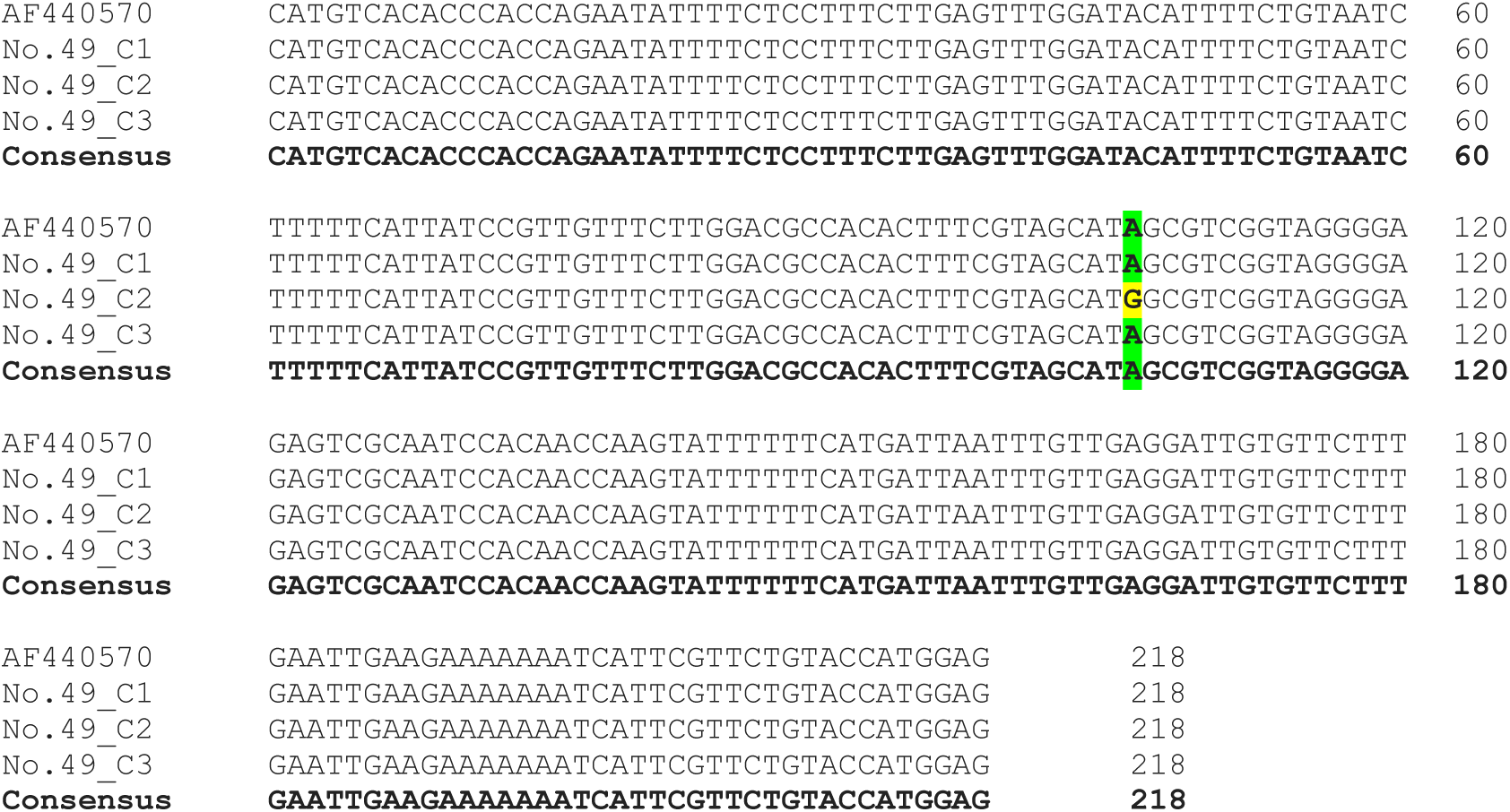

